# Pulsed stimuli entrain p53 to synchronize single cells and modulate cell-fate determination

**DOI:** 10.1101/2023.10.24.563786

**Authors:** Harish Venkatachalapathy, Zhilin Yang, Samira M. Azarin, Casim A. Sarkar, Eric Batchelor

**Author notes:** Correspondence (E.B.); (C.A.S.).

## Abstract

Entrainment to an external stimulus enables a synchronized oscillatory response across a population of cells, increasing coherent responses by reducing cell-to-cell heterogeneity. It is unclear whether the property of entrainability extends to systems where responses are intrinsic to the individual cell, rather than dependent on coherence across a population of cells. Using a combination of mathematical modeling, time-lapse fluorescence microscopy, and single-cell tracking, we demonstrated that p53 oscillations triggered by DNA double-strand breaks (DSBs) can be entrained with a periodic damage stimulus, despite such synchrony not known to function in effective DNA damage responses. Surprisingly, p53 oscillations were experimentally entrained over a wider range of DSB frequencies than predicted by an established computational model for the system. We determined that recapitulating the increased range of entrainment frequencies required, non-intuitively, a less robust oscillator and wider steady-state valley on the energy landscape. Further, we show that p53 entrainment can lead to altered expression dynamics of downstream targets responsible for cell fate in a manner dependent on target mRNA stability. Overall, this study demonstrates that entrainment can occur in a biological oscillator despite the apparent lack of an evolutionary advantage conferred through synchronized responses and highlights the potential of externally entraining p53 dynamics to reduce cellular variability and synchronize cell-fate responses for therapeutic outcomes.

## Introduction

Biomolecular oscillations are observed in a variety of cellular processes such as MAPK signaling^1^, cell division^2^, inflammation^3^, and DNA damage^4^. Dynamical features of these oscillations, such as the amplitude and frequency, encode information about a stimulus and determine the cellular response^5^. However, biological noise can cause variations in both these features, leading to cell-to-cell variations in the downstream response^1,6,7^. Entrainment to an external stimulus has evolved as a strategy to facilitate a synchronized population response in the face of such variability.^8^ This synchrony in cellular response can be crucial for cells in a multicellular organism to respond coherently to regulatory cues. Notably, circadian oscillations drive synchronous periodic behavior in response to 24-hour light-dark cycles across multiple systems within an organism, allowing for coherent metabolic and hormonal dynamics.^9,11,13^ Similarly, oscillations in glycolytic networks^10^ and NF-κB^3,12^ have been shown to respond synchronously under periodic stimuli. This is also typically bolstered by cell-cell communication in the form of paracrine or cell-cell junction-mediated signals.^8,14,15^ However, it is unclear whether this extends to oscillatory contexts where the cellular response is intrinsic to a single cell and such coherence in the response of single cells is not required for effective function.

One such context is oscillations in the p53 transcription factor that occur in response to double stranded DNA breaks (DSBs) induced by stressors such as endogenous replicative stress and γ-irradiation. Here, p53 levels increase and decrease in an oscillatory pattern with an average period of ∼5.5 h in mammalian cells and drive DNA damage responses including DNA repair.^16,17^ There is significant variability in pulse frequency between individual cells as well as time between pulses in an individual cell^18^, potentially due to the intrinsic lack of synchronizing cues in this context in the form of cell-cell communication or a periodic external driver. However, the number of DSBs continues to decrease exponentially in the overall population, apparently robust to these fluctuations.^19,20^ Further, despite cell-to-cell variations in p53 dynamics driving heterogeneous tumor cell behavior under cancer therapies^21–26^, the potential to synchronize these dynamics, to our knowledge, remains unexplored. Therefore, we sought to determine if periodic DSB induction could improve synchrony in p53 oscillations.

Using a mathematical model of the p53 response to NCS-induced DSBs developed by Mönke et al.^19^, we predicted the existence of entrainment in an interval around the natural p53 pulsing period. We validated the prediction experimentally using live-cell fluorescence microscopy and single-cell tracking, demonstrating increased synchronization of p53 dynamics in response to a periodic damage signal. Surprisingly, we experimentally observed entrainment across a wider range of stimuli periods than was computationally predicted. Using parameter sensitivity analysis, we found non-intuitively that the increased entrainment range required a modified p53 DSB response model with a weaker oscillator that would be more susceptible to biological noise. Finally, we showed that the change in p53 pulsing period due to entrainment translates to alteration in downstream target dynamics for cell-fate regulatory genes and such frequency modulation can result in differential cell fate specification.

## Results

### The p53 response is entrainable through periodic DSB induction

To identify whether entrainment of p53 pulses through periodic DSB induction was theoretically feasible (**Figure 1A**), we used a mathematical model of the p53 DSB response calibrated using MCF-7 breast cancer cells^19^ (**Figure 1B**) to simulate repeated DSB induction through the radiomimetic drug NCS and analyzed the subsequent p53 response. From deterministic simulations using an NCS-induced birth rate of *b_s_* = 200 breaks/h, we found a small region around the natural frequency of the system (∼1/5.5 h^−1^) where p53 had the same frequency as the input DSB waveform (phase-locked) (**Figure 1C**). We validated this by calculating the entrainment score^12^, which is defined as the fraction of total power on the Fourier spectrum that corresponds to the input frequency and its harmonics. Since an entrained system would have the same frequency as the input, its corresponding frequency spectrum is expected to show high power at the input frequency and harmonics and low power elsewhere, leading to an entrainment score close to 1. Indeed, this was close to 1 for the regions in which the system was phase-locked, further validating our hypothesis that p53 can be entrained by repeated DSB induction (**Figure 1C**). This did not change significantly as a function of *b_s_* over a range of 50 – 1000 breaks/h (**Figure S1**). Additionally, under noisy DSB dynamics, the entrainment score of an ensemble of 1,000 simulations per input frequency closely followed that of the deterministic system (**Figure 1C**), implying DSB stochasticity was likely not a significant source of variability in entrainment in the conditions analyzed.

**Figure 1:**
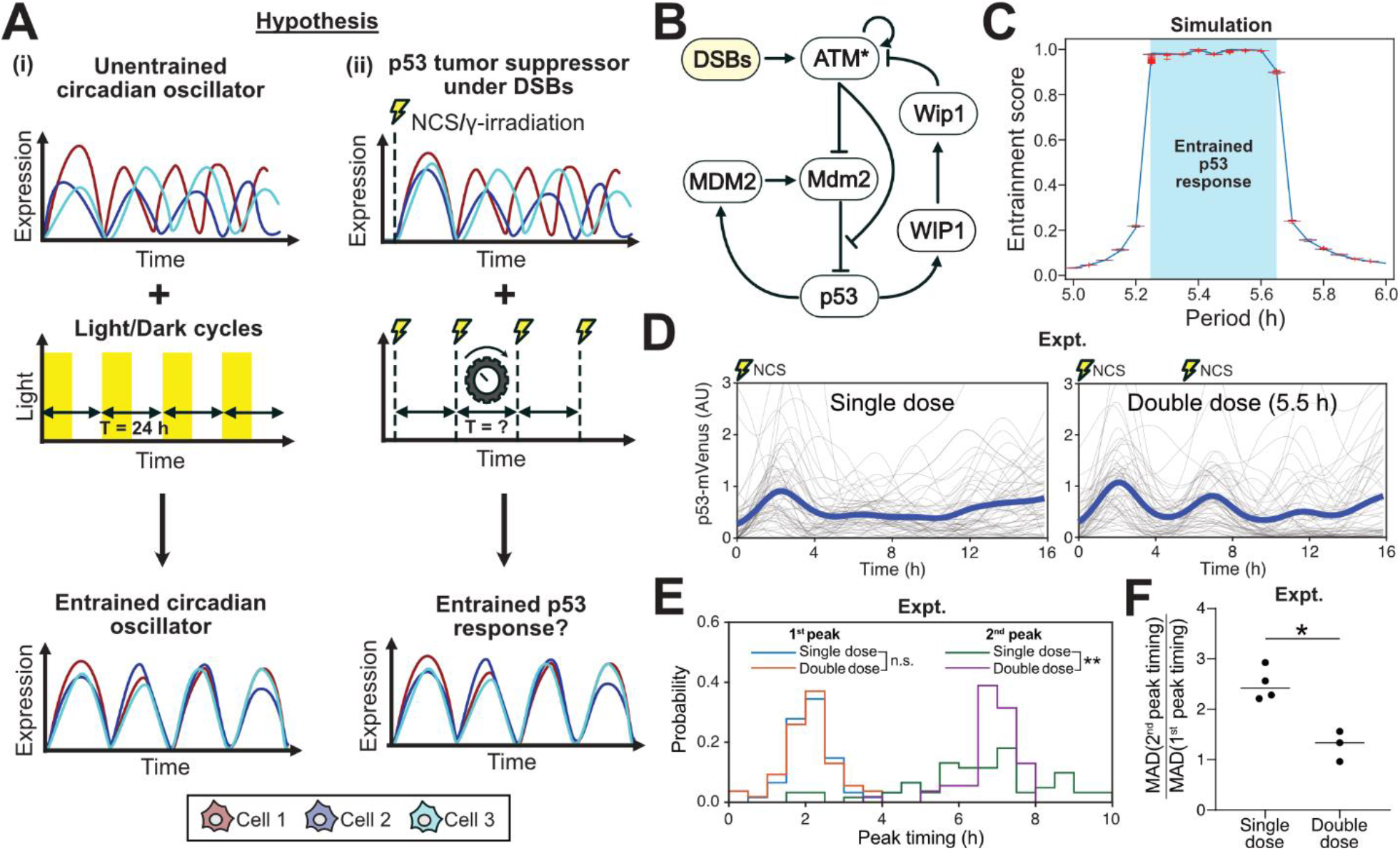
p53 DSB response is entrainable. **(A)** Schematic showing asynchronous oscillations in expression levels of a (i) circadian oscillator that are synchronized under periodic light/day cycles. (ii) We hypothesized that asynchronous p53 oscillations under DSBs could be similarly entrained and synchronized under periodic DSB induction. **(B)** Diagram of the p53 regulatory network under DSBs.^19^ Yellow boxes denote species with stochasticity. **(C)** Entrainment score of deterministic simulations driven by either deterministic (blue line) or stochastic (box plots, 1000 realizations) DSB dynamics. Phase-locked regions in deterministic systems indicated in cyan. **(D)** Single-cell traces (gray) and mean (dark blue) p53-mVenus transgene expression under a single dose of NCS (400 ng/mL) or two doses spaced 5.5 h apart. **(E)** Probability distributions of first and second peak timing. (* p < 0.05; ** p < 0.01; Kolmogorov-Smirnov test) **(F)** Ratio of median absolute deviation (MAD) in peak timing of the second peak to the first, measured across four and three biological replicates for the single dose and double dose conditions, respectively. Each biological replicate consists of at least 45 individually tracked cells. Statistical analysis presented here is from an ANOVA followed by multiple comparisons that was performed on the dataset presented in **Figure S2**.

To validate the computationally predicted conditions of entrainment, we quantified p53 dynamics in p53-mVenus expressing MCF-7 cells that were treated with either a single dose of 400 ng/mL NCS or two doses spaced 5.5 h apart (**Figure 1D**). Both conditions showed a synchronized first pulse of p53 accumulation after treatment, with no statistically significant differences in peak timing distributions (**Figure 1E**). However, the cells that received a single dose of NCS lost synchrony by the second p53 pulse, while cells that received a second dose of NCS at 5.5 h had a significantly narrower second peak timing distribution (p < 0.01) (**Figure 1E**). This was further validated by the smaller increases in the median absolute deviation (MAD) of second peak timing over first peak timing (single dose median: 2.41, double dose median: 1.33; p < 0.05) (**Figure 1F**). Overall, cells that had been treated twice with NCS better retained p53 peak timing synchrony.

### p53 pulses can be entrained over a wider range of stimulus periods than predicted

To experimentally validate the computationally predicted range of stimulus periods across which entrainment is possible, hereafter referred to as entrainability, we treated the MCF-7 p53-mVenus cell line with either a single dose of 400 ng/mL of NCS or two doses spaced 3.0 h – 7.0 h apart, with a resolution of 0.5 h (**Figure 2A, Figure S2**). In contrast to model predictions of a narrow entrainment range of 5.25 h to 5.65 h, we observed statistically significant increases in synchrony in conditions with two doses spaced 4.0 h to 5.5 h apart when compared to a single dose (**Figure 2B**). We did not observe similar synchronization in the 3.0 h, 3.5 h, 6.0 h, 6.5 h, and 7.0 h conditions. Interestingly, the 3.0 h and 3.5 h conditions did not exhibit a synchronous first pulse as typically expected. Instead, we observed cells with either two small pulses spaced close together or one big pulse spanning the approximate duration of the small pulses, thereby exacerbating heterogeneity in the p53 response in comparison to a single dose (**Figure S2**). In contrast, the 6.0 h, 6.5 h, and 7.0 h conditions exhibited a synchronized first pulse similar to a single dose of NCS but were followed by a less synchronized second pulse suggesting loss of entrainment. Surprisingly, inclusion of stochastic mRNA dynamics in computational simulations, a feature shown to widen the entrainment range in other systems^3,12,27^, did not improve entrainability in our p53 model (**Figure S3)**. Taken together, these results indicate that the p53 DSB response is more entrainable than predicted and that the improved entrainability is likely not driven by intrinsic noise.

**Figure 2:**
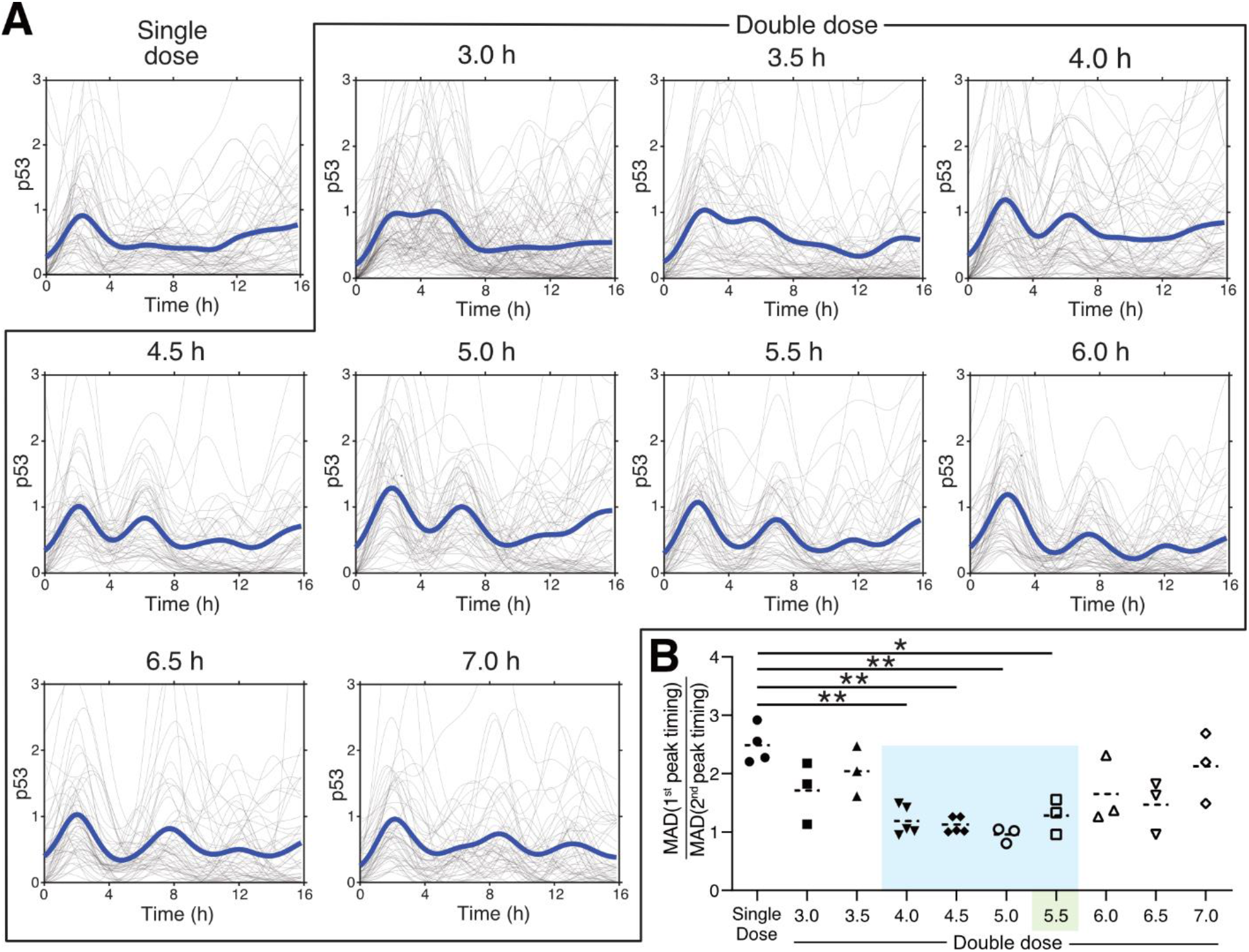
p53 is experimentally entrainable across a wider range of stimulus frequencies than computationally predicted. **(A)** Single cell traces (gray) and mean (dark blue) p53-mVenus expression under a single dose of NCS (400 ng/mL) or two doses spaced 3.0 h to 6.0 h apart. Traces representative of a minimum of three biological replicates; each biological replicate consists of at least 45 individually tracked cells. **(B)** Fold-change in median absolute deviation (MAD) in peak timing from the first peak to the second, measured across a minimum of three biological replicates (* p < 0.05; ** p < 0.01; ANOVA followed by multiple comparisons). Single dose and 5.5 h double dose data same as **Figures 1, S2**.

### Parameter sensitivity analysis identifies oscillation strength as a predictor of entrainment range

Given that intrinsic noise alone was not explanatory of the entrainment range observed experimentally, we performed a parameter sensitivity analysis to examine whether changes in system parameters could explain the experimentally observed range (**Figure 3A**).

**Figure 3:**
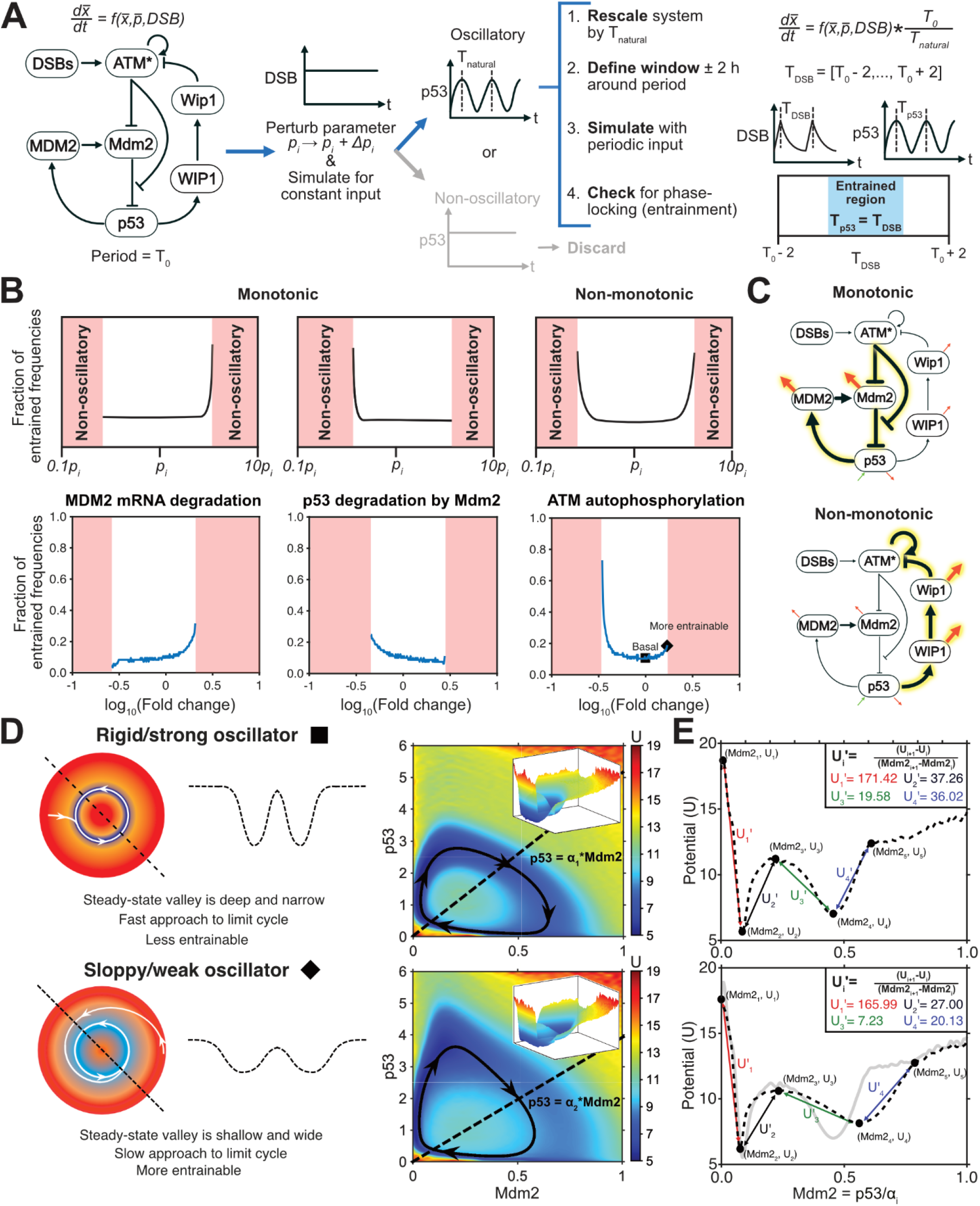
Parameter sensitivity analysis identifies proximity to non-oscillatory regimes and limit cycle rigidity as a positive predictor of entrainment range. **(A)** Schematic depicting the parameter sensitivity analysis workflow. **(B)** Depictions of the two types of entrainment range responses to parameter changes in the system with examples of each type. **(C)** Schematic of the p53 DSB response model with specific interactions that exhibit monotonic or non-monotonic responses highlighted and enlarged in size. Black arrows depict interactions while green and red arrows represent basal production and degradation, respectively. **(D)** Graphical depictions and the underlying energy landscapes of a more rigid, less entrainable oscillator (basal system; black square in ATM autophosphorylation rate diagram in (B)) and less rigid, more entrainable oscillator (ATM autophosphorylation rate changed to 1.67 times the basal value; black diamond in ATM autophosphorylation rate diagram). Rigidity refers to the strength of the limit cycle attractor. Black line represents the deterministic path (limit of zero noise) and arrows represent direction of movement. Dashed line represents the line passing through the origin and the inner maximum of the steady-state valley described by Mdm2 = α_i_*p53 where α_1_ = 4.76 and α_2_ = 4.24. Energy landscapes generated using p53 DSB model with stochastic DSB and mRNA dynamics and system size of 100. **(E)** Slice of energy landscapes corresponding to (D) along the dashed line depicting the change in the rigidity of the steady state. Gray line in the bottom panel corresponds to the more rigid landscape (dashed line in the top panel).

We observed two classes of behavior based on whether the range of input frequencies conferring entrainment changed monotonically or non-monotonically with the parameter. The monotonic class consisted of parameters associated with the p53-Mdm2 loop (e.g., MDM2 mRNA degradation rate and Mdm2 mediated p53 degradation rate). The non-monotonic class consisted of parameters associated with the p53-Wip1-ATM* interactions (e.g., ATM autophosphorylation rate) (**Figure 3B, C**). While some parameters had a stronger effect on entrainability than others (**Figure S4**), the most striking feature was a sharp increase in entrainment range at the cusp of the transition to a non-oscillatory regime in some cases, implying that less robust oscillatory behavior improved entrainment. That is, the range of entrainable frequencies was largest in parameter regimes where small changes in parameters such as those arising from extrinsic noise, a part of natural biological variability, led to the loss of oscillatory behavior. While it is non-intuitive that entrainment, a feature associated with oscillations, correlated with less robust oscillations, we hypothesized that an endogenous oscillator with a weaker limit cycle attractor would be more susceptible to perturbations and thereby more entrainable by an external cue. To validate this, we used a trajectory-based energy landscape methodology^28,29^ to generate the underlying energy landscapes for the basal system and compared it to a more entrainable system (1.67 times the basal ATM autoactivation rate). We expected oscillators with weaker (less rigid) limit cycle attractors to have a shallower stable steady-state valley that is reflective of higher susceptibility to external perturbations while oscillators with stronger (more rigid) limit cycle attractors would have a deeper valley (less affected by external forces) (**Figure 4D**). By comparing the two landscapes, we found that the valley around the mean system path in the more entrainable system was wider and had a gentler gradient around it (**Figure 4E**). This was further corroborated by examining the potential across a line connecting the origin and the inner maximum within the circular steady-state valley. This showed higher stable steady-state potentials and lower slopes around the states in the more entrainable landscape, two characteristics that suggest greater responsiveness to external cues. Taken together, the results validate our hypothesis that a less rigid oscillatory system is more affected by external perturbations and thus is more entrainable and that the p53 DSB response needs to be underpinned by such a weak oscillator to recapitulate our experimental observations.

**Figure 4:**
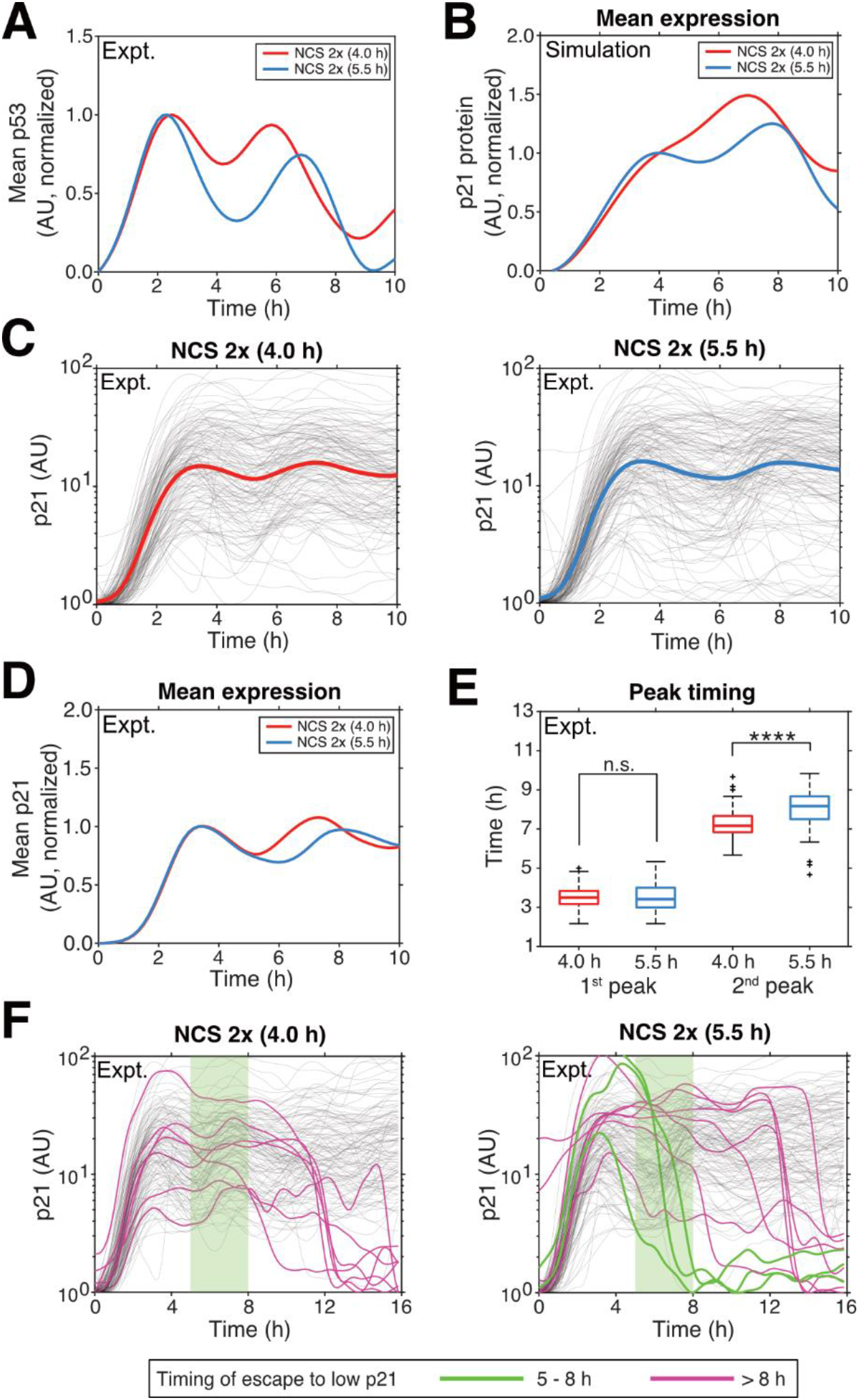
Entrainment of p53 pulses modulates downstream p21 expression. **(A)** Mean p53-mVenus expression under two doses of NCS spaced 4.0 h (red) or 5.5 h (blue) apart, rescaled from 0 to 1 such that the first peak for both curves are the same height. **(B)** Model prediction of downstream p21 expression driven by p53 expression shown in (A). **(C)** Single cell traces (gray) and mean (red and blue lines) p21-mVenus expression of responding cells under the conditions in (A) with n = 135 and n = 148 for the 4.0 h and 5.5 h conditions, respectively obtained from three biological replicates. **(D)** Mean p21 expression for the two conditions shown in (C). Experimentally observed curves are comparable to computationally predicted curves in (B). **(E)** Comparison of p21 peak timings for the first and second peaks in the 4.0 h and 5.5 h conditions showing no differences for the first peak, but a later second peak for the 5.5 h condition. Comparisons were performed using Wilcoxon rank sum test (**** p < 0.0001). **(F)** Traces of p21 expression in NCS 4.0 h and NCS 5.5 h conditions demonstrating escape to a low p21 state either between 5.0 h to 8.0 h (green) or after 8.0 h (magenta). Traces filtered for escapers prior to 5.0 h. Light green highlight represents the time period from 5.0 h to 8.0 h.

### p53 pulse modulation through entrainment changes downstream mRNA target dynamics in a half-life-dependent manner

As p53 dynamics control cell fate under DSBs, we hypothesized that changes in p53 pulsing frequency due to entrainment alter the downstream response. We validated this by examining the expression of the cell-cycle inhibitor p21, a canonical p53 downstream target. Comparing the mean p53 expression level across single cells for NCS stimulus periods of 4.0 hours and 5.5 hours, we observed a shift in the peak timing of the second p53 pulse (**Figure 4A**). Using a mathematical model of p21 expression^7^, we predicted that these p53 dynamics result in higher expression of p21 at the population level between 5.0 h and 8.0 h as well as an earlier second peak in the NCS 4.0 h condition (**Figure 4B**). To validate this, we quantified p21 protein dynamics in MCF-7 cells tagged with mVenus at the p21 C-terminal genomic locus under different NCS treatment intervals (**Figure 4C**). We only considered cells that had a minimum normalized expression of 10^0.5^ units at t < 3.0 h to filter out cell-cycle dependent delays in p21 upregulation^30^. Consistent with the model, we found higher mean p21 expression in the NCS 4.0 h condition between 5.0 h to 8.0 h compared to the NCS 5.5 h condition (**Figure 4D**). Further, we found that the second peak timing of p21 expression was later for cells administered a second dose of NCS at 5.5 h compared with 4.0 h (p < 0.0001; Wilcoxon rank sum test). No differences were observed in the first peak timings between the two conditions (**Figure 4E**). Finally, based on a study by Reyes et al.^7^, we predicted that lower expression of p21 in the NCS 5.5 h condition that we had observed from 5.0 h to 8.0 h compared to the NCS 4.0 h condition can result in stochastic transitions to a low p21 state during this time, characteristic of escape from cell cycle arrest. Indeed, while there are cells that revert to a low p21 state in both the NCS 4.0 h and NCS 5.5 h conditions (magenta traces, **Figure 4F**), escape from this state between 5.0 h to 8.0 h occurs only in the NCS 5.5 h condition (green traces, **Figure 4F**).

We also observed increased mRNA expression of *CDKN1A* (p21 mRNA) as well as other p53 targets (*GADD45A, TRIAP1,* and *FAS*) under administration of a second dose of NCS at 4.0 h compared with 5.5 h between 5.0 h and 8.0 h (**Figure S6**). This effect was most prominent and consistent in *GADD45A*, but less so in the other genes, appearing in only one or two out of three biological replicates. This may be due to differences in mRNA stability – *GADD45A* is highly unstable and follows p53 dynamics more closely in comparison to more stable species such as *CDKN1A*, *TRIAP1,* and *FAS*.^31^ Therefore, any changes in p53 dynamics would be better reflected in *GADD45A* dynamics as opposed to more stable downstream mRNA targets. Indeed, mathematical modeling of mRNA transcription using p53 dynamics from **Figure 4A** as an input, we find that less stable mRNA targets show higher differences in expression between the 4.0 h and 5.5 h NCS interval treatments (**Figure S7**). Overall, this suggests that entrainment of the single-cell p53 response can modulate downstream target expression in a target stability-dependent manner.

## Discussion

Periodic external cues can synchronize biological rhythms across cells. This is particularly important for driving a coherent response in metabolic regulation as well as paracrine/endocrine signaling in multicellular organisms. ^8,9,11,12^ In this study, we demonstrated that this phenomenon also extends to the p53 oscillatory response to DSBs, a context where cell-to-cell coherence does not underpin an effective cellular response as measured by successful DNA repair. By periodically inducing DSBs through NCS (a small molecule γ-irradiation analog), cell-to-cell heterogeneity in the period of p53 pulses was reduced significantly compared to a single initial dose. Interestingly, the range of entrainment observed experimentally (> 1.5 h) (**Figure 2**) was significantly larger than computationally predicted (∼0.5 h) even under high DNA damage (**Figure 1C, Figure S1**). Further, in contrast to other systems where intrinsic noise widens the entrainment range^3,12,27^, there was no such intrinsic noise-driven entrainment range widening in the p53 system (**Figure S3**). Instead, using parameter sensitivity analysis, we found that the widened entrainment range in the p53 system is due to the presence of a less rigid endogenous oscillator than predicted by the mathematical model (**Figure 3**). This phenomenon has also been observed in the circadian system.^32,33^

Energy landscape analysis of this phenomenon showed that a landscape which permitted more uniform cell behavior through entrainment was characterized by a wider stable steady state valley under a constant input, a feature associated with more heterogeneity. In other words, a system that exhibits noisier oscillations under a constant input is expected to be more entrainable and better synchronized under periodic inputs. While this seems counterintuitive, it is consistent with the expected behavior of a weak attractor. Such an attractor permits significant noise-driven deviations from the mean under steady state (a key feature of p53 oscillations under DSBs^7,16^) while simultaneously being more malleable to an external cue, allowing for a more uniform response under external perturbations. Taken together, these observations suggest that p53 dynamics under DSBs are governed by a weak oscillator and that this feature is responsible for better entrainability of the system. However, it is unclear whether this can be simply recapitulated by reparameterization of the model or if it requires the addition of missing regulatory interactions. For example, incorporating bidirectional regulation of DSB repair and p53 could lead to better entrainment if there is a rigidity mismatch between these two interconnected oscillators.^34^

Finally, we found that controlling p53 dynamics through entrainment enabled modulation of the expression of downstream targets of p53, including targets involved in cell fate determination. When examining p21 expression in single cells (**Figure 4**), we found that a higher entraining frequency resulted in greater mean p21 expression across the population, which enabled better maintenance of cell cycle arrest compared with a lower entraining frequency. However, the effects on other p53 targets were less clear. Out of four downstream targets examined (**Figure S6**), only the least stable species *GADD45A* reproducibly displayed altered expression dynamics at the population level. The effect of entraining treatments was qualitatively, but not statistically, reproducible in the more stable targets including *CDKN1A* (p21 mRNA), which exhibited stimulus frequency-dependent dynamics at the single-cell level. Mathematical modeling attributed this lack of statistical significance to a diminished difference in mRNA dynamics between the two entraining frequencies for more stable targets (**Figure S7**), which were likely more difficult to recapitulate experimentally due to natural biological variability, especially with bulk measurements of a non-homogeneous population. Overall, these findings imply that less stable p53 targets, such as those involved with DNA repair and cell cycle arrest^31,35,36^, are most likely to be affected by entrainment.

In comparison to other biological oscillators such as the circadian rhythm and NF-κB, the inherent value of synchronized p53 oscillations is less clear. It is possible that entrainment is simply a consequence of the evolution of oscillations in p53; this dynamic behavior has increased information transfer capacity in comparison to single-value steady states under noise and can therefore enable cells to respond sensitively to DNA damage while maintaining the potential to recover^5,7^. Alternatively, given the effects of p53 on circadian rhythm regulation^37^, metabolism^38^, and NF-κB^39,40^, it is possible that synchrony in p53 oscillations shapes more effective cellular responses in these contexts despite not being required for DNA repair. Examining the interplay of these oscillatory systems and p53 in regulating gene expression and cell fate determination could help elucidate evolutionary advantages of entrainability in p53 oscillations.

From an engineering perspective, the increase in cell-to-cell synchrony in p53 dynamics observed here offers an avenue for designing cancer therapies that can result in more effective tumor cell clearance. Strategies involving entrainment can mitigate effects such as fractional survival of tumors by providing a more homogenous initial population that is expected to respond more uniformly to a secondary therapy^41^. Further, the ability to modulate downstream gene expression through stimulus frequency can be used to bias cellular response towards specific cell fates – for example, stimulating p53 at frequencies that result in increased DNA repair gene expression and better maintenance of cell cycle arrest can aid in better recovery of cells in cases such as damage due to mustard gas exposure.^45^

Overall, this study demonstrates the entrainability of p53 oscillations under DSBs, elucidating fundamental oscillator characteristics that underpin this phenomenon. This entrainment also led to stimulus frequency-dependent modulation of downstream target genes responsible for cell fate determination. Future work focused on utilizing the reduced single-cell heterogeneity as well as differential gene expression demonstrated here may aid in the design of more effective therapeutic strategies in contexts where p53 modulates cellular behavior such as cancer and chemical damage remediation.

## Materials and Methods

### Mathematical modeling of p53 dynamics

Deterministic simulations were carried out in MATLAB (Mathworks, Natick, MA, USA), and stochastic simulations were carried out using the Gillespie algorithm32 in C++. A mathematical model of the p53 response under double stranded DNA breaks (DSBs) developed by Mönke et al.^19^ was used in all simulations, with modifications as specified. The mRNA synthesis rates of MDM2 and WIP1 were increased by 20% each (from 1.0 AU/h to 1.2 AU/h) to reduce the observed p53 pulse frequency from once every 7.0 h to once every 5.5 h, matching the expected period in MCF-7 cells.^16^ Other parameters could also be potentially varied to get this same effect. We chose MDM2 and WIP1 production rates based on the sensitivity analysis by Mönke et al.^22^ Additionally, the functional form of DSB sensing by ATM, *f(DSB)* (**Figure S8A**), was changed from a saturable equation, *f(DSB) = DSB/(γ + DSB)*, to a non-saturable one, *f(DSB) = log(DSB/γ + 1)*, to reflect the shift from oscillatory to sustained p53 dynamics observed experimentally under high DNA damage.^4^ Both of these functions show very similar behavior for small numbers of DSBs (**Figure S8B**). However, the saturable function approaches a maximum of 1 and continues to drive oscillatory behavior for large numbers of DSBs (**Figure S8C**). In contrast, the non-saturable log function causes a shift from oscillatory to sustained p53 dynamics under high damage (**Figure S8D**). Values for all rate constants used are provided in **Table S1**.

#### Deterministic model

The double stranded break and repair process from the original model^19^ was changed from a discrete stochastic birth-death process to a deterministic equation: *d(DSB)/dt = b – r*DSB*. Following Mönke et al.^19^, for the first hour after NCS treatment, the birth rate was changed to *b_s_*, a parameter proportional to the NCS concentration. After one hour until further damage was inflicted, the birth rate reverted to a basal birth rate *b_b_* = 2.3 breaks/h. Additionally, DSBs were repaired with a constant death rate *r* = 0.315 h^−1^ throughout the simulation. For repeated treatment, this resulted in a sawtooth-like waveform where there was nearly linear accumulation of DSBs with a slope proportional to *b_s_* for 1 hour followed by exponential decay (**Figure 7E**). Simulation files are provided in **File S1**.

#### Stochastic model

The model included two key sources of noise: (1) DSB birth-death process, and (2) mRNA production and degradation. Due to its independence from the rest of the model, noise due to stochastic DSB generation and repair was implemented by separately generating the appropriate waveform using the Gillespie method^42^, and then feeding this waveform as a piecewise function of time into the time-stepping algorithm. The mean of the DSB process realizations closely followed the sawtooth-like deterministic shape (**Figure S8F**). The model with DSB noise was simulated in MATLAB R2020a using the *ode23s* function with the DSB waveform input as a piece-wise function such that for *t_i_ < t < t_i+1_*, *DSB = DSB_i_*, where *DSB_i_* is the number of double-strand breaks as per the stochastic birth-death model. To include noise from the stochastic production and degradation of mRNA species (WIP1 and MDM2), we implemented a hybrid stochastic-deterministic simulation in C++. In this implementation, the equations for WIP1 and MDM2 mRNAs were solved using the Gillespie method and all other equations were solved deterministically using discrete time steps between each time step of the Gillespie algorithm. The number of time steps was chosen to have a smaller average time step duration than the DSB noise only simulation from MATLAB. The number of DSBs were updated according to the pre-generated waveform such that *DSB = DSB_i_* when *t > t_i_*, *DSB = DSB_i+1_* when *t > t_i+1_*, and so on. Each stochastic simulation was run 1,000 times, as determined by the convergence in entrainment score (**Figure S8G**). Simulation files are provided in **File S1**.

#### Entrainment characterization

For deterministic simulations, the system was considered phase-locked and therefore entrained if, at steady state, the period was within 0.1% of the input period (an order of magnitude lower than the period grid size/interval when evaluating entrainment). Due to variability in the period of oscillation in stochastic systems, this approach was not applied there. Instead, the degree of entrainment was calculated using the entrainment score (*E*), following the procedure used by Gupta et al.^12^ Briefly, this score is the fraction of power at the input frequency and its harmonics, compared to the entire frequency spectrum as calculated by a one-sided discrete Fourier transform (DFT, custom script in MATLAB). The DFT calculates power *P* at a set of frequencies *f* dependent on trajectory length and discretization. Due to finite observation time, we will have a finite number of frequencies which may or may not include the exact input frequency and the corresponding harmonics. Instead, the spectral power of the system will be spread around the components of the DFT that are near the exact frequencies. In consideration of this numerical issue, we used a frequency window *w*, that was 1% of the width of the natural frequency of the system *f_natural_* within which the corresponding power for each frequency was calculated. The entrainment score was then calculated as follows,

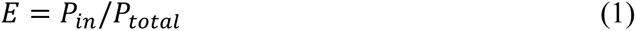

where,

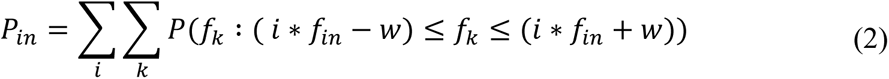

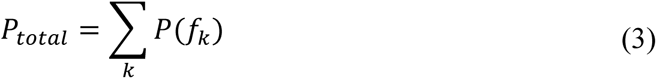

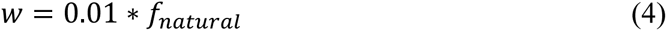

*f_k_* is the k^th^ frequency of the DFT, *i* is an integer corresponding to the *i^th^* harmonic, *f_in_* is the input frequency*, w* is the window size around the frequencies of interest, and *f_natural_* is the natural frequency of the system.

All simulations were run for 5,000 hours for entrainment characterization. As opposed to shorter times, this duration exhibited minimal fluctuation in the entrainment score in phase-locked regions (**Figure S8H**). These fluctuations are artifacts caused by the finite numerical nature of the DFT, as explained in the previous paragraph.

#### Parameter analysis for system entrainment

##### Overview

Parameters were varied, one at a time, and the resulting system was classified as oscillatory or non-oscillatory based on the total Fourier power of the normalized waveform. The system was then rescaled by multiplication with 5.5 (natural period of the basal system) and division with the natural period of the modified system. Entrainment, as defined by phase-locking, was examined in a window of ± 2 h around the natural pulsing frequency of the system.

##### Parameter variation

Simulations of the fully deterministic system were used to characterize the effect of individual parameters on entrainment. Each parameter was separately varied in a 10-fold range below and above the default value in increments of 0.005 on the log_10_ scale.

##### Constant DSB simulation and classification into oscillatory/non-oscillatory

The system was first simulated for a constant DSB input of 200 for *t* = 5,000 h. The p53 trajectory after 1,000 h (in intervals of 0.05) was classified as oscillatory or non-oscillatory based on the total power on the Fourier spectrum. The trajectory was first rescaled linearly from 0 to 1 (*rescale* function in MATLAB) to remove variability in power due to amplitude and then was used to calculate the one-sided DFT. If the total power of the DFT was > 0.2, it was classified as oscillatory. This threshold was determined by simulating 1,000 Latin-hypercube sampled parameters in a 2-fold range, for constant DSB inputs from 0 to 1,000 in increments of 20, and examining the distribution of total power (**Figure S8I**). A clearly bimodal distribution arose: one population of trajectories had a minimum power of 0.2, likely representing oscillatory cases, while the other population had a maximum power of ∼0.1, corresponding to non-oscillatory cases. A few randomly selected trajectories from both parts of the distribution were visually inspected to validate this. Non-oscillatory cases were discarded while oscillatory cases moved on to the next stage.

##### System rescaling with natural period

Oscillatory systems were then rescaled by multiplying the right-hand side of the differential equation expressions by 5.5 h (natural period of the basal system) and dividing it by the natural period of the oscillatory system (in h). This ensures commensurate comparisons of entrainment ranges across different parameter sets where the system may have different natural periods.

##### Entrainment curve generation and characterization

The natural period of the oscillatory trajectory was determined by the most prominent peak on the DFT. A window (± 2 h, in increments of 0.05 h) was defined around this period and entrainment was characterized for periodic inputs by examining phase-locking within this region. The effect of each parameter on entrainment was examined by the number of points entrained for each parameter value divided by the total number of points in the window.

### Mathematical modeling of downstream targets

p21 protein dynamics in **Figure 4B** were simulated using a mathematical model of p21 mRNA and protein dynamics developed by Reyes et al.^7^ All rate constants were the same as the original publication except for *K*_0_(Michaelis constant of p53-dependent p21 transcription) which was set to 0.67. This was done to account for the differences in the scaling of p53 traces between this work and Reyes et al.^7^ General mRNA dynamics in **Figure S7** were obtained using a standard p53-dependent transcription model with p53 levels from **Figure 4A** rescaled from 1 to 5 as the input, consistent with Porter et al.^31^ Rate constant values were the same as the original work. Simulation files are provided in **File S1**.

### Cell culture

MCF-7 cells transfected with p53-mVenus and H2B-Cerulean were maintained in RPMI-1640 media supplemented with 10% (v/v) FBS and 1% (v/v) Pen-Strep (all reagents from Life Technologies) in a humidified incubator at 37 °C and 5% CO_2_. MCF-7 cells with p21 tagged fluorescently at the endogenous genomic locus (a kind gift from Sabrina L. Spencer^43^) were maintained in the same medium. Due to the loss of DHB-mCherry (Cdk2 sensor) and/or H2B-mTurquoise (nuclear marker) in subpopulations of these cells, fluorescence-activated cell sorting (FACS) using a BD FACSAria II instrument was used to obtain cells that retained expression of both constructs.

### Fluorescent clonal cell line generation for live-cell imaging of p53 expression

Lentiviral particles containing pLentiPGK-Hygro-DEST-H2B-mCerulean3 (kind gift from Markus Covert^44^; Addgene: 90234) vector were produced in HEK293T (ATCC) cells using Lenti-X Packaging Single Shots (TakaraBio) using the manufacturer’s protocol. The resulting particles were used to infect MCF-7 cells expressing a previously characterized p53-mVenus fusion.^4^ Cells stably expressing both constructs were selected in cell culture media containing 50 μg/mL hygromycin and 400 μg/mL neomycin (Life Technologies). Clonal cell lines were generated by limiting dilution. Clones were then screened for the presence of Cerulean fluorescence. Additionally, the cells were validated to exhibit the previously reported pulsatile p53-mVenus expression dynamics under 400 ng/mL of neocarzinostatin (NCS; MilliporeSigma).

### Live-cell imaging and single-cell tracking

For live-cell imaging, cells were plated on 96-well No. 1.5 glass-bottom dishes (Mattek) to be 60% confluent at the start of treatment. Prior to imaging, cells were washed with DPBS (1x), after which transparent RPMI-1640 media (Life Technologies) – lacking phenol red, riboflavin, and L-glutamine, and supplemented with 2% (v/v) FBS and 1% (v/v) Pen-Strep – was added. Cells were imaged with a Nikon Eclipse Ti2 inverted fluorescence microscope equipped with an automated stage (Prior) and a custom chamber to maintain 37 °C, 5% CO_2_, and high humidity. Images were collected every 10 minutes for the brightfield, YFP, and CFP channels using a 20x CFI Plan Apochromat Lambda (NA=0.75) objective (Nikon). Exposure time for each channel was set to be <500 ms. The ND2 format imaging data were exported as individual tiff files for each channel, time point, and stage position using Bio-Formats^46^ command line tools. Cells were tracked semi-automatically in p53Cinema^7^ using the nuclear marker H2B-mCerulean in the CFP channel, agnostic to p53-mVenus expression in the YFP channel. p53-mVenus expression for each cell track was then obtained using the getDatasetTraces_fillLineageInformation function with a sampling radius of 5 pixels applied to the YFP channel.

### p53 single-cell trace processing and analysis

All p53-mVenus traces for each stage position were first divided by the mean p53-mVenus fluorescence reading at the start of the experiment (*t* = 0) at that position to account for illumination variability between stage positions. Any missing readings were replaced by the p53-mVenus expression at the previous time point within that trace (*fillmissing* function in MATLAB). The traces were then smoothed using a gaussian moving filter (*smoothdata* function in MATLAB) with a window size corresponding to a real-world time of 3 hours and 20 minutes; this window size aids in smoothing out smaller noisy fluctuations while retaining overall pulsatility of the system. The final traces were obtained by subtracting the minimum value of each trace from itself providing a lower bound at 0 for the p53 signal in each cell. The analysis script is provided in **File S1**.

### Peak detection and peak-timing analysis

Peaks were detected for each trace and the mean of all traces using the *findpeaks* function in MATLAB and specifying a set number of peaks to be detected. To quantify the peak-timing variability for each n^th^ peak, we first identified the peak in each trace that is closest to the n^th^ peak of the mean p53 signal and then calculate the median absolute deviation in peak-timing in this set. Using this method instead of simply considering the n^th^ peak for each trace avoids erroneous propagation of confounding effects due to cells that did not pulse during the first dose, but rather pulsed starting from subsequent doses, while still accounting for whether a cell pulsed or not. The analysis script is provided in **File S1**.

### Gene expression quantification

Cells were plated on 35 mm dishes to be 60% confluent at the start of treatment. To obtain cell pellets, each dish was harvested at the specified time point by scraping in DPBS followed by centrifugation and aspiration of the supernatant. The resulting pellets were immediately stored at −80 °C. Total RNA was extracted from cell pellets using a RNeasy Mini Kit (QIAGEN). Lysis was performed using 350 µL of the provided lysis buffer and the lysate was passed through a QIAshredder column (QIAGEN). RNA from the lysate was obtained by following the manufacturer’s protocol and quantified using a Nanodrop 2000 (ThermoFisher Scientific). cDNA was prepared by loading 2 µg of RNA per sample into a 20 µL reverse transcription (RT) reaction using a High-Capacity cDNA Reverse Transcription Kit (ThermoFisher Scientific) as per the manufacturer’s protocol. The resulting cDNA mixture was diluted 1:5 (20 µL of cDNA added to 80 µL of nuclease-free water) before usage. A 20 µL reaction mix was prepared using 10 µL iTaq Universal SYBR Green Supermix (Bio-Rad), 1 µL 10 µM forward/reverse primer mix (sequences from Porter et al.^31^), 8 µL nuclease-free water, and 1 µL template. qRT-PCR was then performed using a CFX Connect Real-Time PCR Detection System (Bio-Rad) with the following conditions: hot start (95 °C for 2 min), and 40 cycles of PCR (96 °C for 5 s, 64 °C for 20 s), ending in melt curve acquisition (64 °C–95 °C with 0.5 °C resolution). Melt curves (Change in relative fluorescence units per unit temperature vs. Temperature) were visually inspected for existence of a single peak to validate specific amplification for each gene. The threshold cycle count values (*Cq*) for each gene were obtained using the Bio-Rad CFX Manager 3.1 software. The fold change in expression was calculated by 2^−ΔΔ*C*_*q*_(*X*,*t*)^ where ΔΔ*C*_*q*_(*X*, *t*) is defined as:

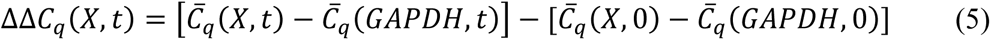

where **C̄*_q_* (*X*, *t*) is the mean *Cq* value for gene *X* at time *t*.

The error in fold-change was obtained by propagating the variability in *Cq* values involved in the calculation. The error in gene X at time t was obtained as follows:

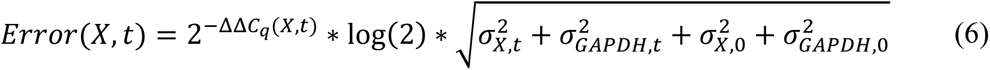

where 2^−ΔΔ*C*_*q*_(*X*,*t*)^ is the fold change and 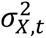 is the variance in *Cq* of gene *X* at time *t*.

### Statistical analysis

All statistical analyses were carried out using GraphPad Prism 9.0 or MATLAB. Experimental data in **Figure 1** and **Figure 2** passed a normality check using the Shapiro-Wilk test (**Figure S8J**) and were compared using one-way ANOVA followed by multiple comparison testing (Prism). Fold changes in variability for NCS 4.0 h (n = 6, originally) and 4.5 h (n = 7, originally) were run through an outlier detection test (Prism), and one and two data point(s) for each condition, respectively, were identified as outliers and discarded prior to further statistical analysis. qPCR data in **Figure 4C** were analyzed using two-way ANOVA followed by multiple comparison testing (Prism). Peak timings in **Figure 4E** failed a normality check by using the Kolmogorov-Smirnov test and were therefore compared using a non-parametric Wilcoxon rank sum test (*kstest* and *ranksum* functions in MATLAB). qPCR data in **Figure S6** were analyzed on the *Cq* scale. The mean expression of each gene from three biological replicates was analyzed using a repeated measures ANOVA followed by multiple comparisons. Expression of each gene within a biological replicate was also individually analyzed from technical replicates using an ANOVA followed by multiple comparisons.

### Data Availability

Upon acceptance of the manuscript for publication, computational code for all models will be publicly available through GitHub, and imaging datasets will be publicly available through Image Data Resource.

## Acknowledgments

The authors thank Kala Guettler for assistance with single-cell tracking. The authors also thank Sabrina Spencer for providing the p21-Cdk2 tagged cell line used in this study. This work was supported by funding from the University of Minnesota (S.M.A. and E. B.) and the National Institutes of Health (R35GM136309 to C.A.S. and R01GM149666 to E.B.), as well as access to high-performance computing resources from the Minnesota Supercomputing Institute and fluorescence-activated cell sorting services from the University Flow Cytometry Resource at the University of Minnesota.

## Supplementary Material

**Figure S1:**
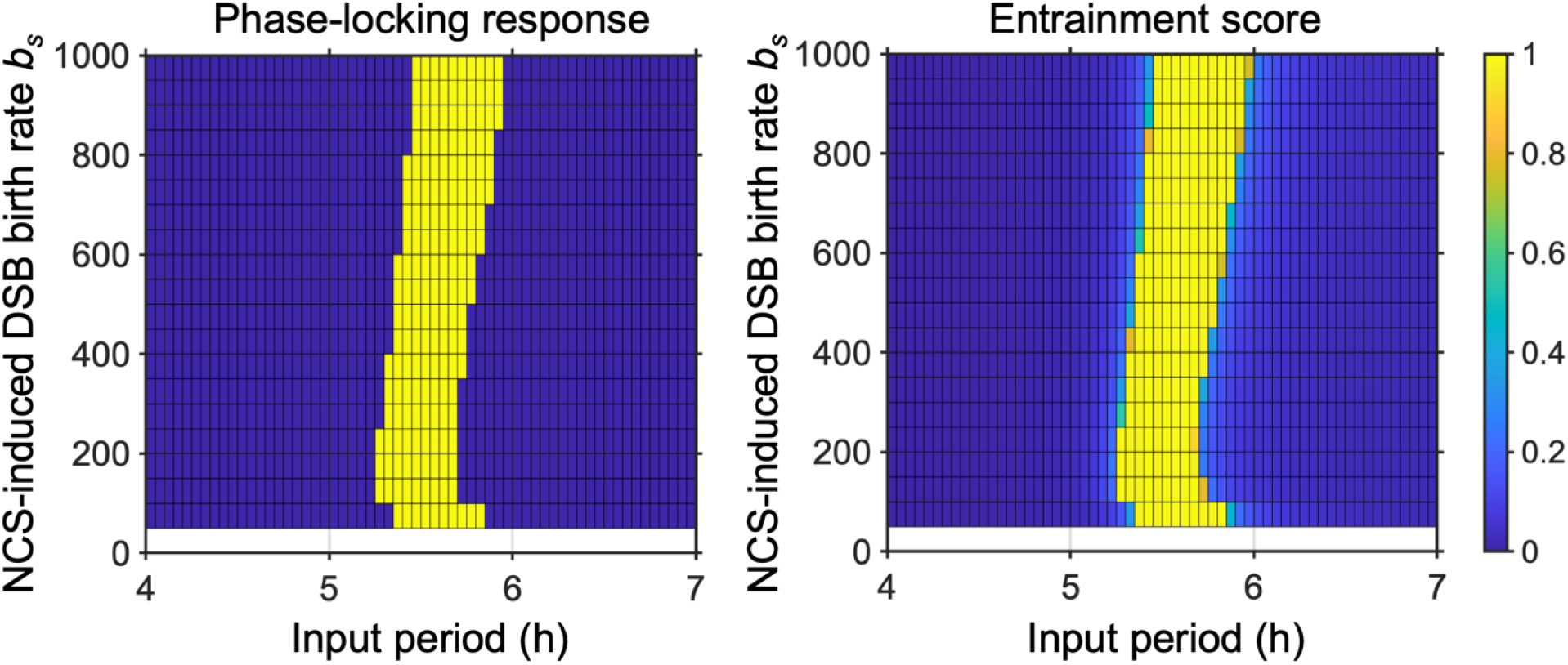
Phase-locking response and entrainment scores as a function of NCS-induced birth rate and input period. Yellow regions represent phase locking while blue represents no phase-locking.

**Figure S2:**
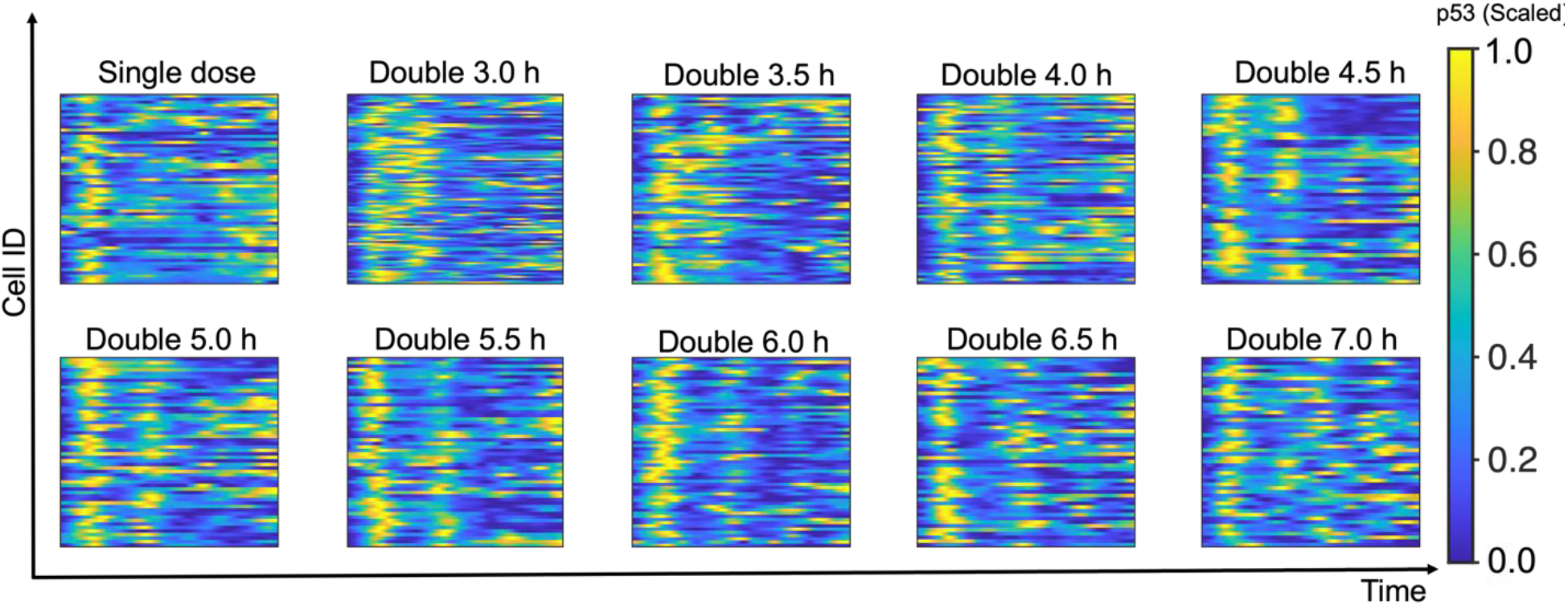
p53 responses from **Figures 1, 2** represented as a heat map diagram where each p53 trace is rescaled from 0 to 1.

**Figure S3:**
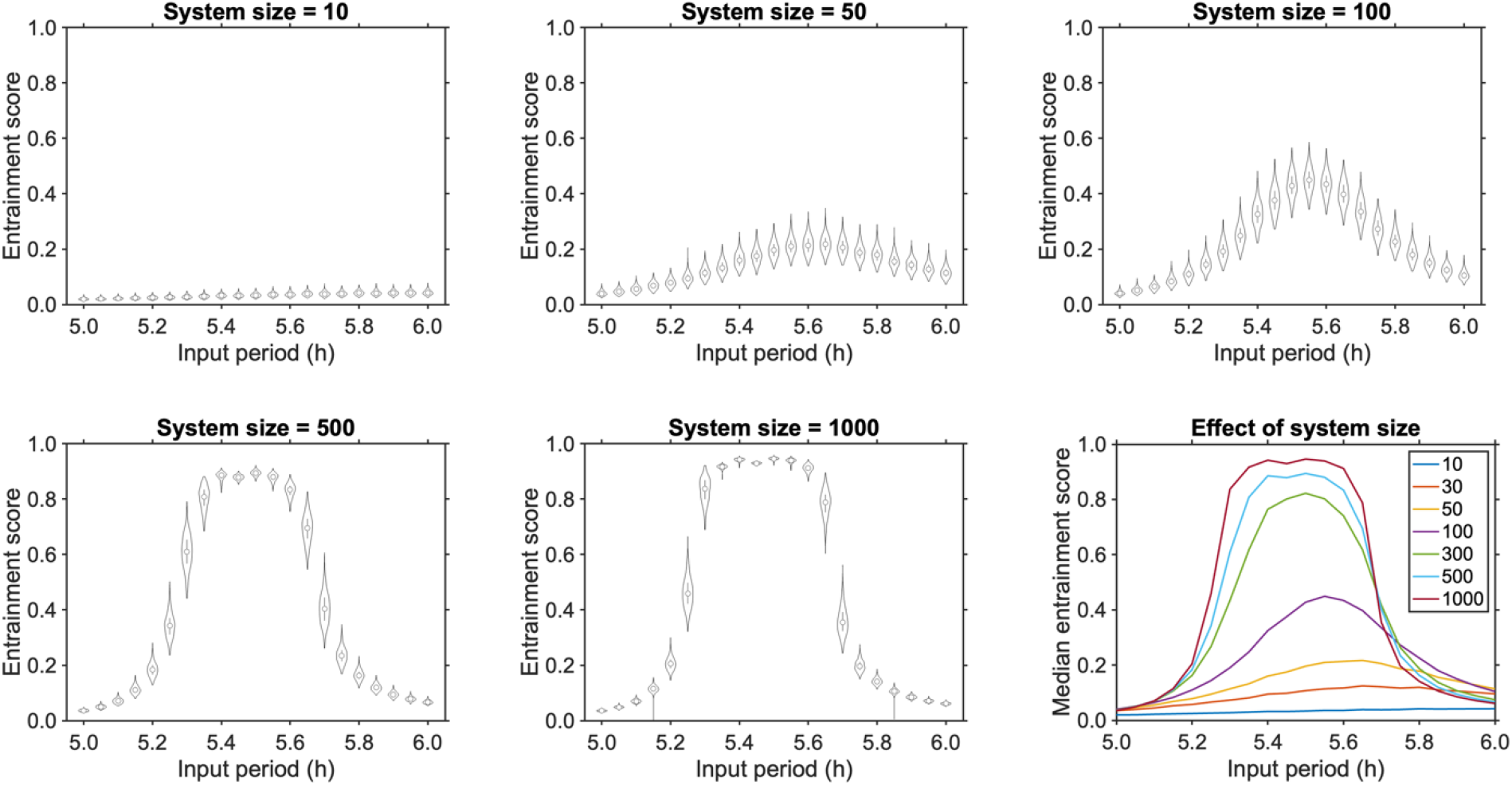
**(A – E)** Violin plots of the top 99% of entrainment scores for 1,000 realizations of the p53 DSB response in different system sizes at different NCS input periods for an NCS-induced break rate of 200 breaks/h. Here, system size refers to the factor used to convert concentration to number of molecules in the stochastic simulations. Larger system size implies more molecules and, therefore, less noise. Conversely, smaller system size implies fewer molecules and more noise. **(F)** Mean entrainment score of 1,000 realizations of the p53 DSB response in different system sizes at different NCS input periods for an NCS-induced break rate of 200 breaks/h. Overall, there is no increase in entrainment range due to intrinsic noise in comparison to the deterministic system.

**Figure S4:**
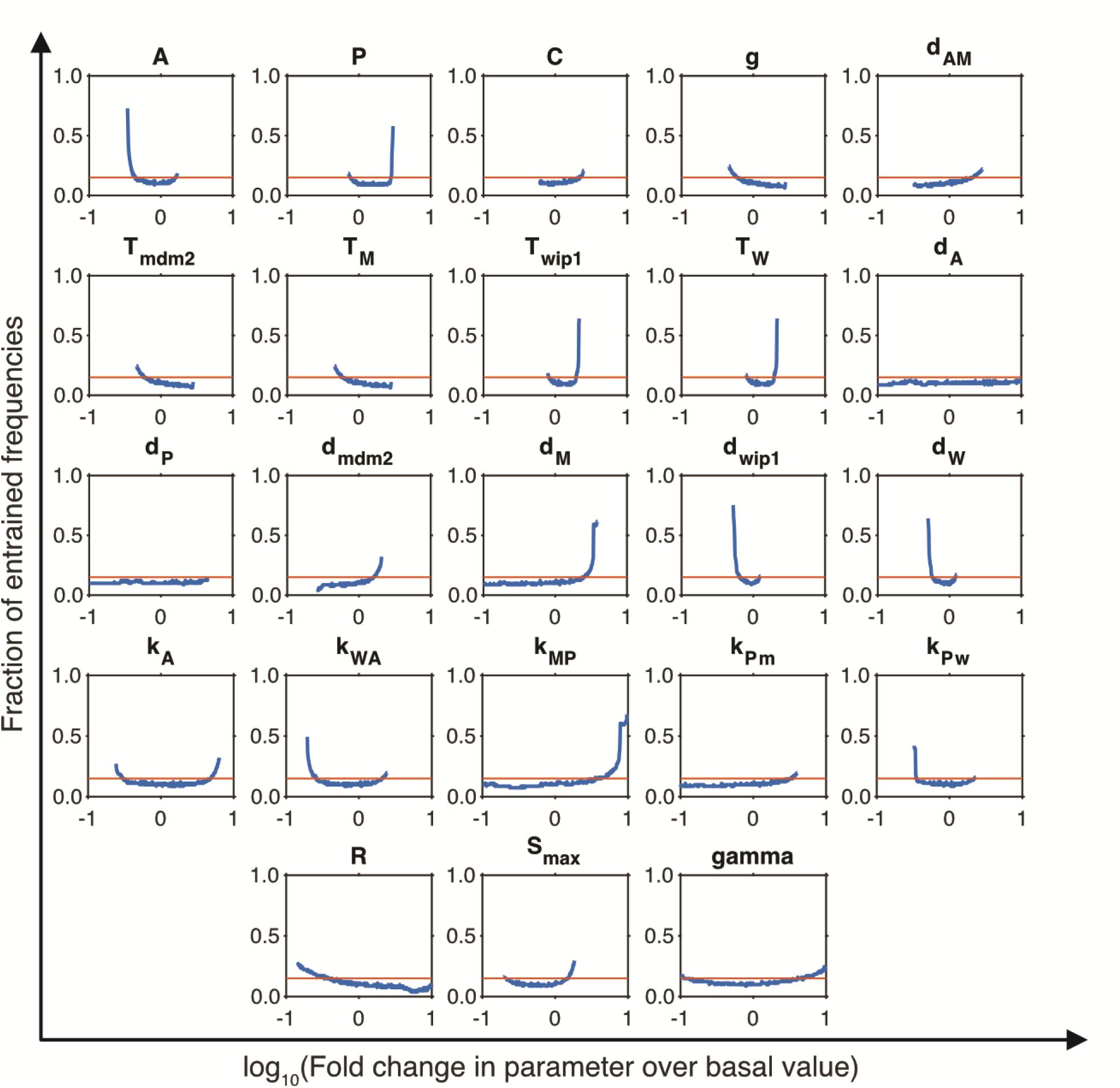
Effect of log_10_ fold-change in each parameter on the fraction of input periods in a scaled window around the natural oscillatory period of the system. Effect of parameters considered significant if the y-axis value exceeded 0.15 (orange line, ∼50% increase over basal entrainment range). Explanation of symbols provided in **Table S1**.

**Figure S5:**
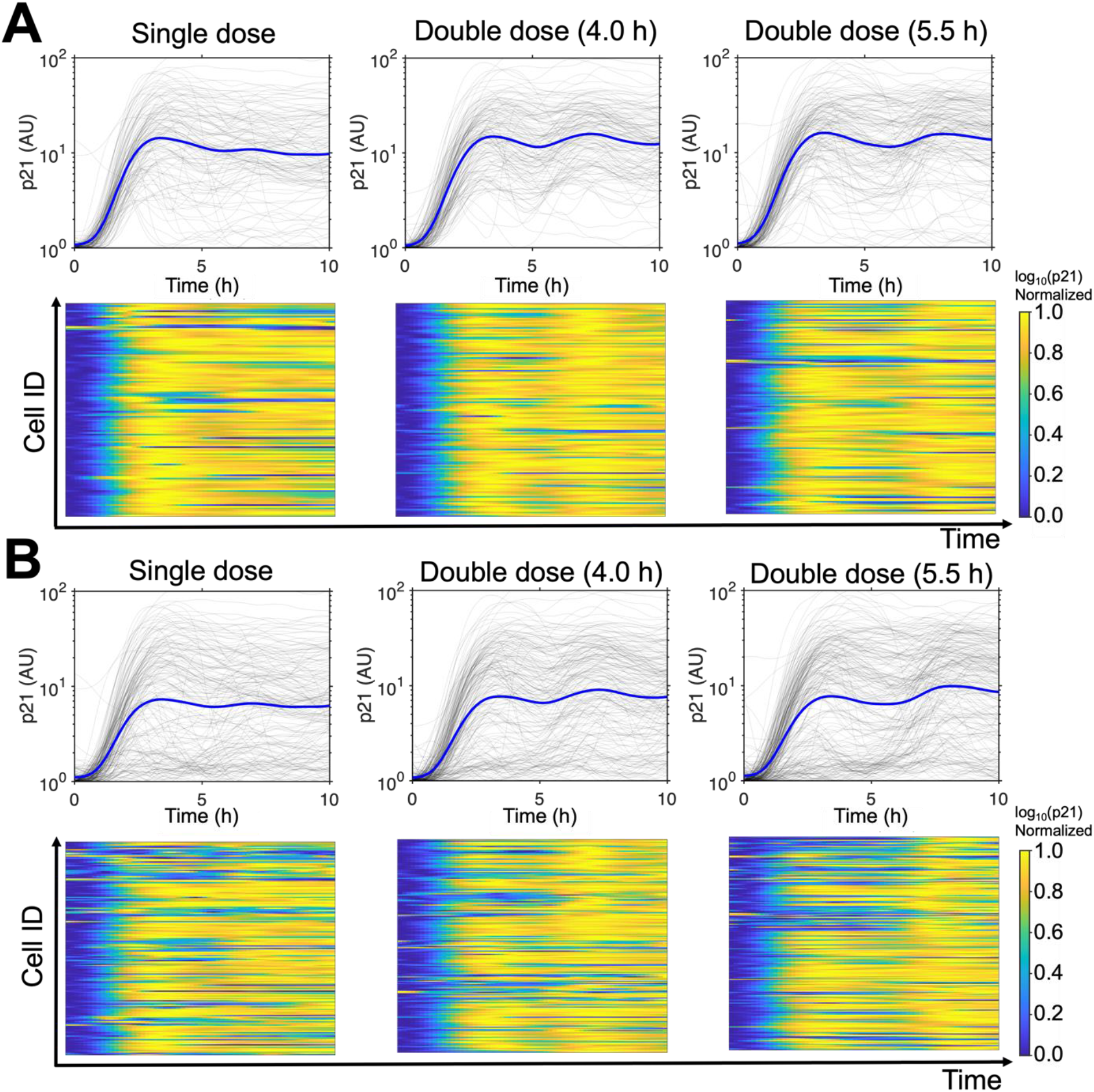
**(A)** Single-cell traces (gray) and mean p21 expression values (blue) for conditions from Figure 4 along with a single dose of NCS as well as heat map diagrams for these conditions where each p21 trace is rescaled from 0 to 1. n = 121, 135, and 148 for the single dose, double dose (4.0 h) and double dose (5.5 h) conditions, respectively **(B)** All single-cell traces of p21 expression without filtering for responding cells showing variability in timing of p21 induction despite the same treatment conditions (as reported previously^30^). n = 175, 190, and 216 for the single dose, double dose (4.0 h) and double dose (5.5 h) conditions, respectively

**Figure S6:**
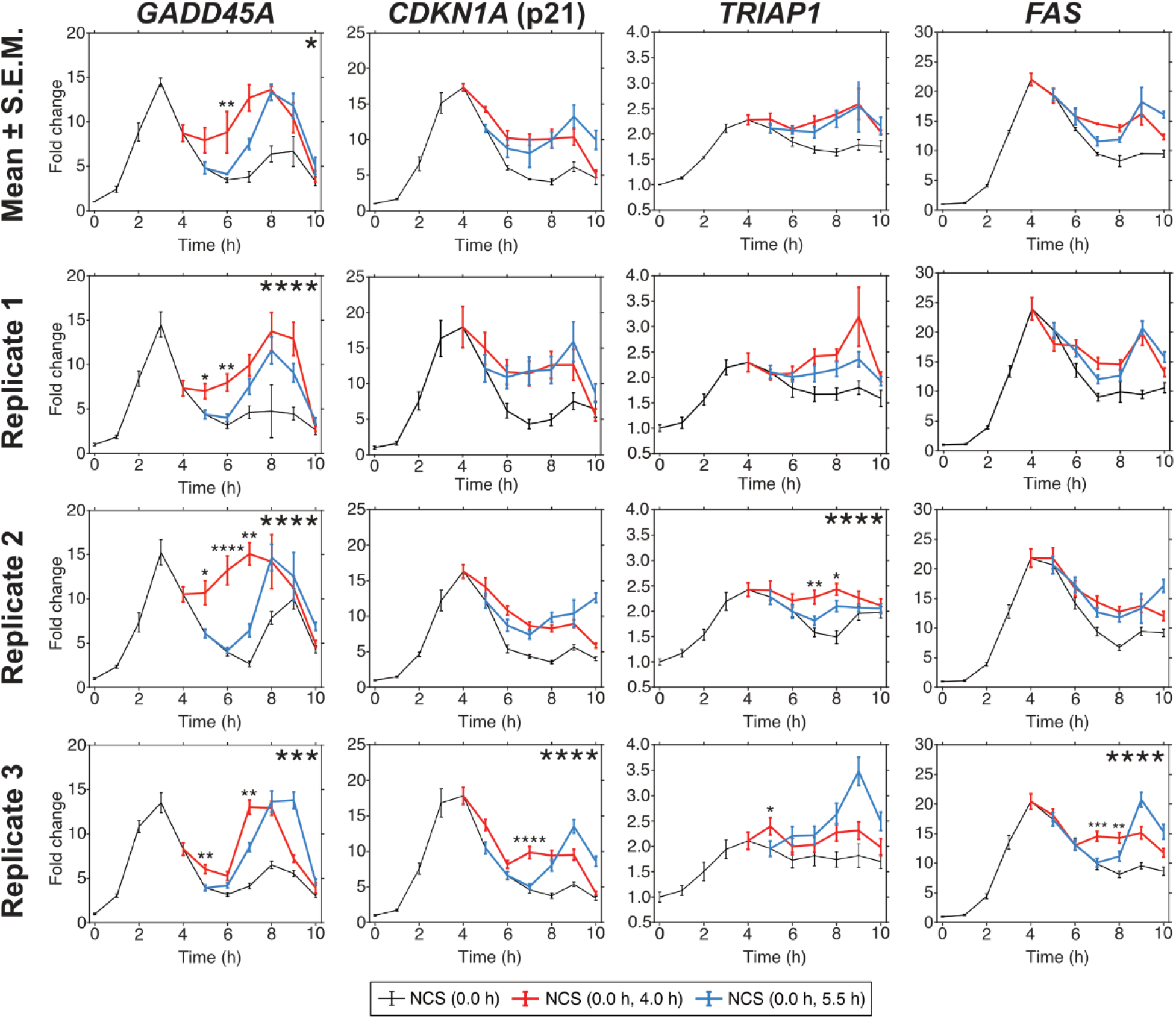
Gene expression of *GADD45A*, *CDKN1A*, *TRIAP1*, and *FAS* as measured by qRT-PCR in MCF-7 p53-mVenus cells treated with a single dose (black lines), or double dose of NCS at 4.0 h or 5.5 h intervals (red or blue lines, respectively). Top row is the mean and SEM of three biological replicates. Bottom rows show the mean and standard deviation of the technical replicates within each biological replicate. Statistical significance of the difference in gene expression between the 4.0 h treatment and 5.5 h treatment from 5.0 h to 8.0 h is shown on the top right of each plot. Statistical significance of the differences in gene expression at a specific time between the two conditions is shown on top of each point. (* p < 0.05; ** p < 0.01; *** p < 0.001; **** p < 0.0001)

**Figure S7:**
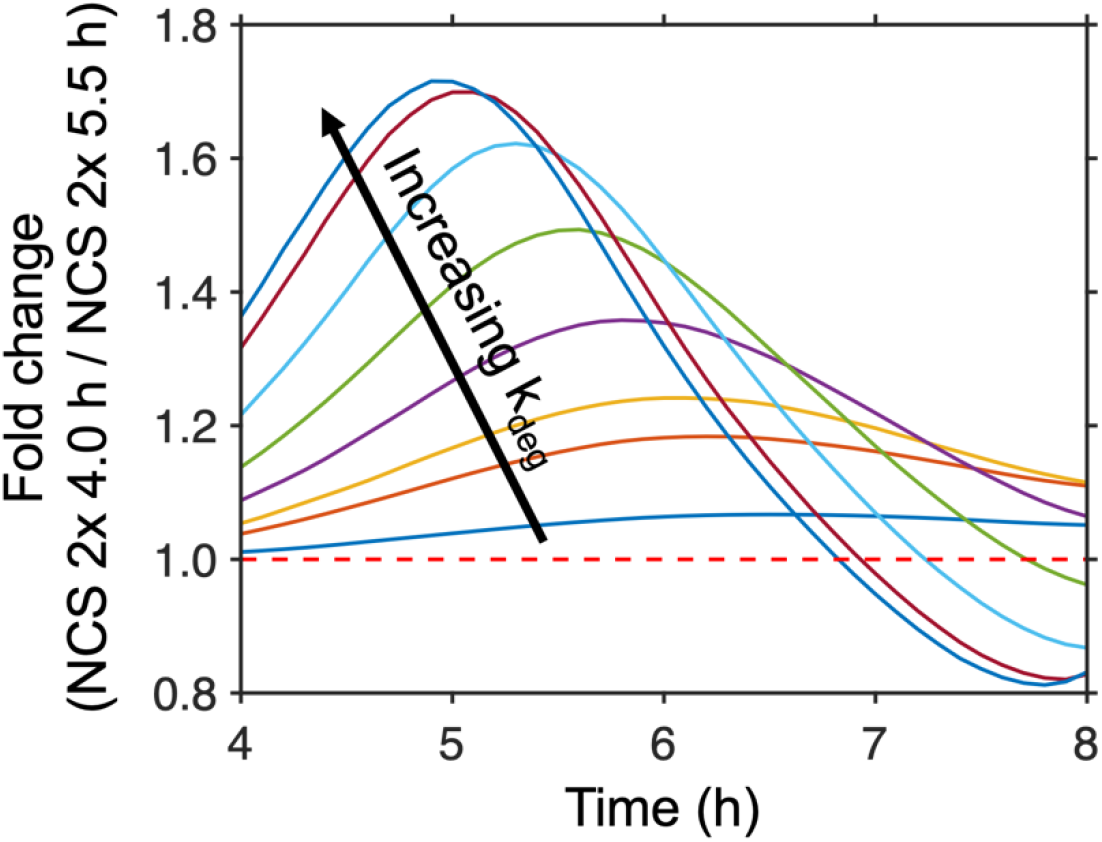
Mathematical modeling prediction of the fold change in mRNA expression with an NCS double dose in a 4.0 h interval compared to a 5.5 h interval for a range of mRNA degradation rates (k_deg_ = 0.01, 0.1, 0.2, 0.5, 1, 2, 5, and 10). Red dashed line represents no change.

**Figure S8:**
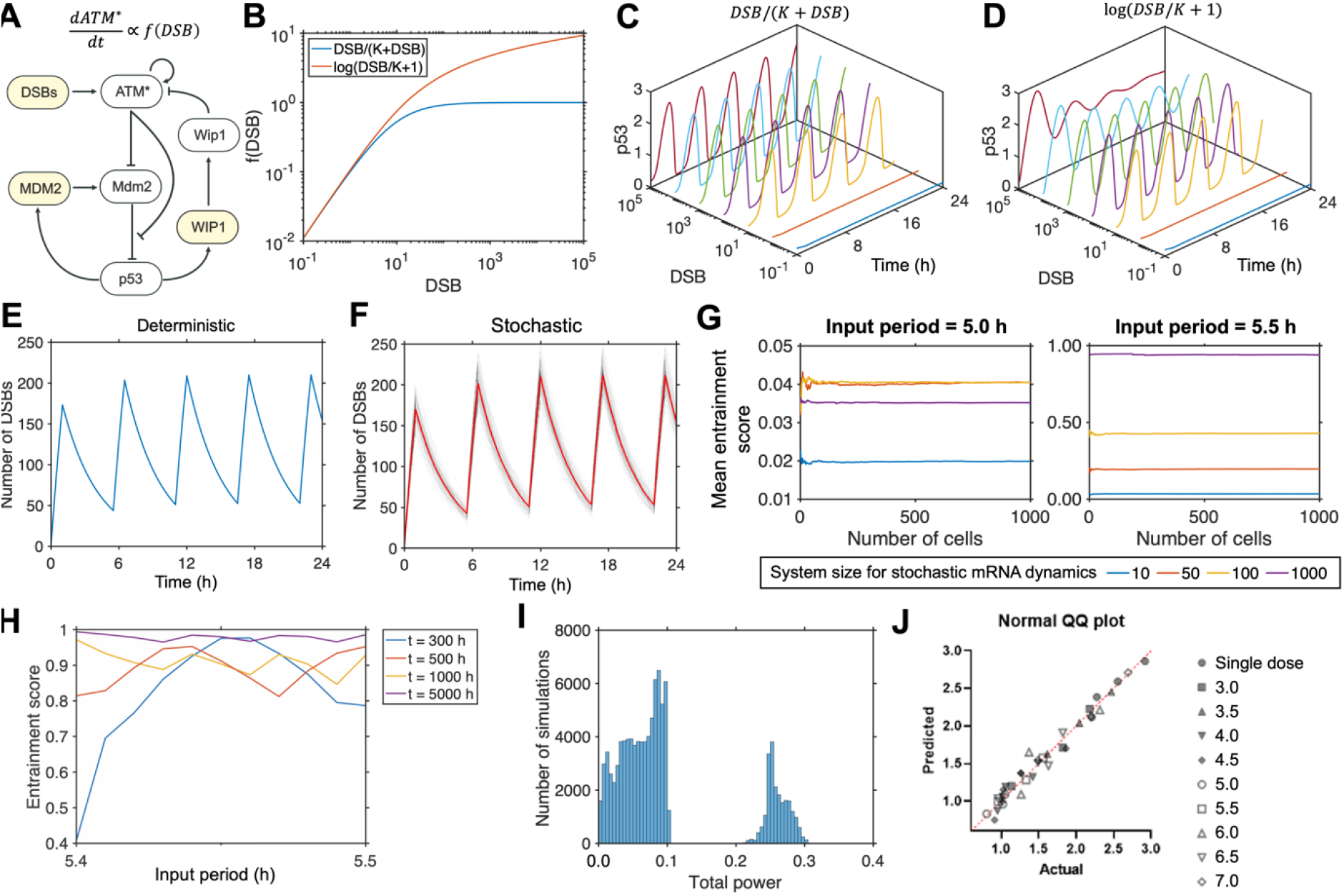
**(A)** Schematic of the p53 DSB response network where the rate of change in ATM* is directly proportional to a DSB input function **(B)** Plots of the saturable and non-saturable DSB input functions for different values of DSBs. **(C, D)** Simulations of the p53 system with the saturable **(C)** and non-saturable **(D)** DSB input functions for different values of DSBs showing similar behavior at low damage but divergent behavior at high damage. **(E)** Deterministic DSB induction and repair with NCS-induced damage being applied every 5.5 h with a NCS-induced birth rate of 200 breaks/h, showing a characteristic sawtooth shape. **(F)** Stochastic DSB inductions and repair with same conditions as (E). Gray lines represent individual realizations of the stochastic process; the red line represents the mean across multiple runs. **(G)** Plot showing the average entrainment score as a function of the number of stochastic realizations for a non-entraining and an entraining input period under different levels of noise. Each plot shows convergence well before 1,000 instances. **(H)** Entrainment score of the fully deterministic system for different simulation lengths showing fluctuations in shorter simulations due to the finite numerical nature of the DFT used to calculate the entrainment score. **(I)** Histogram of the total spectral power at steady state for systems with parameter sets randomly sampled in a 2-fold range around the basal values and simulated with 0 to 1,000 DSBs, in intervals of 20, as the input. **(J)** QQ plot of the Shapiro-Wilk normality test for data in **Figures 1, 2.**

**Table S1.** Parameter symbols and descriptions.

| **Symbol** | **Parameter description** | **Value** |
| --- | --- | --- |
| A | ATM autophosphorylation rate | 30.5 |
| P | Wip1-dependent ATM dephosphorylation rate | 22 |
| C | Basal p53 synthesis rate | 1.4 |
| g | Mdm2-dependent p53 degradation rate | 2.5 |
| d<sub>AM</sub> | ATM-dependent Mdm2 degradation rate | 20 |
| T<sub>mdm2</sub> | MDM2 mRNA production rate | 1.2 |
| T<sub>M</sub> | Mdm2 protein production rate | 4 |
| T<sub>wip1</sub> | WIP1 mRNA production rate | 1.2 |
| T<sub>W</sub> | Wip1 protein production rate | 1 |
| d<sub>A</sub> | Basal ATM dephosphorylation rate | 0.16 |
| d<sub>P</sub> | Basal p53 degradation rate | 0.1 |
| d<sub>mdm2</sub> | MDM2 mRNA degradation rate | 1 |
| d<sub>M</sub> | Mdm2 protein degradation rate | 2 |
| d<sub>wip1</sub> | WIP1 mRNA degradation rate | 1.3 |
| d<sub>W</sub> | Wip1 protein degradation rate | 2.3 |
| k<sub>A</sub> | Michaelis constant for ATM autophosphorylation | 0.5 |
| k<sub>WA</sub> | Michaelis constant for Wip1-dependent ATM dephosphorylation | 0.14 |
| k<sub>MP</sub> | Michaelis constant for Mdm2-dependent p53 degradation | 0.15 |
| k<sub>Pm</sub> | Michaelis constant for p53-dependent MDM2 mRNA production | 1 |
| k<sub>Pw</sub> | Michaelis constant for p53-dependent WIP1 mRNA production | 1 |
| R | Strength of inhibition of Mdm2-dependent p53 degradation by ATM | 2 |
| S<sub>max</sub> | Strength of ATM activation by DSBs | 0.2 |
| gamma | Michaelis constant of ATM activation by DSBs | 9 |

